# Splice Isoform Induced Selective Inhibition of Vesicular Monoamine Transporter Assembly Revealed by Multi-level Combinatorial Analysis

**DOI:** 10.1101/2025.11.29.691296

**Authors:** Alper Karagöl, Taner Karagöl

**Affiliations:** Istanbul University Istanbul Medical Faculty, Istanbul, Turkey

**Keywords:** Truncated isoforms, Monoamine transporters, mRNA, Synaptic vesicle, Neurotransmission

## Abstract

Synaptic vesicle loading of monoamines represents a rate-limiting step in neurotransmission, and its regulation directly shapes dopaminergic, serotonergic, and noradrenergic signaling. Vesicular Monoamine Transporters (VMATs), key determinants of quantal content and synaptic tone, have traditionally been thought to function primarily through their full-length canonical isoforms. However, alternative splicing generates isoforms that remain largely uncharacterized in membrane transporter biology, despite growing proteogenomic evidence supporting their translation. Here, we reveal a potential mechanism by which a truncated splice isoforms of VMAT1 interferes with canonical oligomer formation. Using a combination of structural docking, atomistic lipid bilayer molecular dynamics, and Poisson-Boltzmann Surface-Area (MMPBSA) binding energy decomposition, we characterize the interactions and co-evolution between the truncated isoform and the canonical protein. While the isoform retains partial interface compatibility, it exhibits higher binding affinity for the canonical VMAT than the canonical homodimer itself. Mechanistically, this molecular latch shifts the rules of engagement from a contest of size (surface area) to one of intensity (charge), effectively resolving the David-versus-Goliath conflict: the system energetically favors the heterodimer, ensuring that the smaller isoform consistently dominates over the full-length protein. Functionally, this positions the truncated isoform as a negative regulator of VMAT oligomerization. These findings redefine VMAT assembly as a splicing-sensitive checkpoint and provide a novel mechanistic framework for understanding synaptic dysfunction in neuropsychiatric and neurodegenerative disorders.

## Introduction

The transport of monoamines (dopamine, norepinephrine and serotonin) is a fundamental aspect of synaptic physiology, directly linked to motor control, mood regulation, and cognitive flexibility [1,2,3,4,5]. Once they are moved intracellularly, monoamines are recycled or they are metabolized by monoamine oxidases [3,4,5,6,7]. By determining the quantal size of monoamine-containing vesicles, vesicular monoamine transporters (VMATs) exert rate-limiting control over synaptic output and directly influence neuropsychiatric vulnerability [1,2,3,4,5]. Given their essential role in maintaining neurotransmitter homeostasis, VMATs has emerged as a pharmacologically critical target in neuroprotection and neurodegenerative disease intervention [1,4,5,8]. Vesicular transport efficiency, transporter turnover and conformational changes could have system-wide consequences on neuropsychiatric disease risk [8,9,10,11].

The vesicular monoamine transporters (VMATs) have long been assumed to function primarily through its full-length isoform. However, transcriptome-wide splicing surveys reveal the truncated VMAT variants, the function of which remains unexplored [12]. Splice variants, though lacking domains specific for transport, could be indicated in disease mechanisms [13,14]. Understanding these interactions is crucial, as they may underlie physiological compensatory mechanisms or pathological interference in neurotransmitter regulation [15,16,17].

Our recent findings have revealed the existence of truncated isoforms derived from alternative splicing within the glutamate transporter locus and these isoforms may interfere with canonical transporter oligomerization, leading to modulatory effects on glutamate transporter assemblies [16,17]. On these isoforms interaction domain is often conserved, leading to interactions with canonical forms despite lacking full molecular architecture [16]. Unlike glutamate transporters in the plasma membrane, VMATs are localized to synaptic vesicles, a vesicular compartment characterized by a chemically distinct and highly curved lipid environment [18,19,20,21,22]. This difference is non-trivial: synaptic vesicles possess a unique lipidomic signature, enriched in cholesterol, phosphatidylserine, and polyunsaturated phospholipids, with asymmetric leaflet composition [20,21,22]. Similarly, despite having similar transport functions, vesicular and plasma glutamate transporters are structurally distinct [23,24]. Building a realistic synaptic vesicle membrane model, therefore, represents a major technical challenge in biomolecular simulations, yet it is essential for capturing lipid-sensitive isoform interference mechanisms [16,25]. To address this, we have reconstructed a chemically accurate synaptic vesicle lipid bilayer and embedded both canonical VMAT homodimers and heteroassemblies containing truncated isoforms.

In this study, we establish a multi-level computational framework that begins with genomics-guided identification of truncated VMAT isoforms derived from alternative splicing. We then performed AlphaFold3 multimer prediction to assess their capacity to engage canonical VMAT protomers at the assembly interface. To capture the vesicular biophysical environment, we constructed fully atomistic synaptic vesicle-mimetic lipid bilayer systems, incorporating realistic lipid composition. By integrating 50ns atomistic molecular dynamics simulations with membrane-adapted MM/PBSA binding free energy decomposition, we quantitatively resolve the stability, energetic penalties, and interface disruption patterns associated with isoform-canonical hetero-oligomer formation [16]. This combinatorial approach allows us to link sequence-level isoform diversity to membrane-dependent assembly energetics, providing a thermodynamically grounded model of isoform-induced inhibition of oligomerization.

Our findings suggest VMAT oligomerization is a pharmacologically actionable target in monoaminergic regulation. We propose that such isoform-driven regulation represents a broader strategy for fine-tuning monoamine storage capacity under conditions of synaptic plasticity, metabolic stress, or pathological transporter dysregulation. By uncovering how alternative splicing intersects with VMAT oligomerization, this study opens a new conceptual framework in which isoform biology and lipid biophysics converge to shape pharmacoresponsiveness and monoaminergic disease susceptibility.

## Results and Discussions

### Structural Bioinformatics Analyses of VMAT Isoforms

The vesicular monoamine transporter (VMAT) family exhibits remarkable structural diversity through alternative splicing, generating isoforms with systematically reduced transmembrane (TM) domain architecture (Supplementary Figures S1,S2,S3,S4,S5,S6). VMAT1 canonical isoform (P54219; 525 aa, 12 TM) maintain the full major facilitator superfamily (MFS) fold comprising pseudo-symmetrical N-terminal (TM1-6) and C-terminal (TM7-12) domains (Fig. 1, Supplementary Figure S1). The 53-residue deletion in P54219-2 isoform (P-isoform) occurs while preserving 11 TM helices (472 aa, 11 TM), indicated by nearly identical biophysical parameters (pI 5.58-5.59, GRAVY 0.583-0.585). In contrast, truncated isoforms display progressive TM domain loss: Q96GL6 (Q-isoform) likely retains only TM1-8 (385 aa, 8 TM), eliminating the entire C-terminal structural module; E5RK12 (258 aa, 4 TM) represents a single structural quarter. These systematic truncations create a hierarchical series of structural modules spanning from complete transporters to minimal membrane-anchored regulatory domains. In Q96GL6, the absence of TM9-12 eliminates the C-terminal helical bundle responsible for completing the MFS symmetry, rendering this isoform structurally incomplete as an independent transporter. However, the preserved TM1-8 region encompasses the entire N-terminal bundle and part of the central hinge. The most truncated variant, E5RK12, retains only four transmembrane helices (TM1-4) corresponding to the N-terminal half of the first MFS bundle.

**Fig 1.**
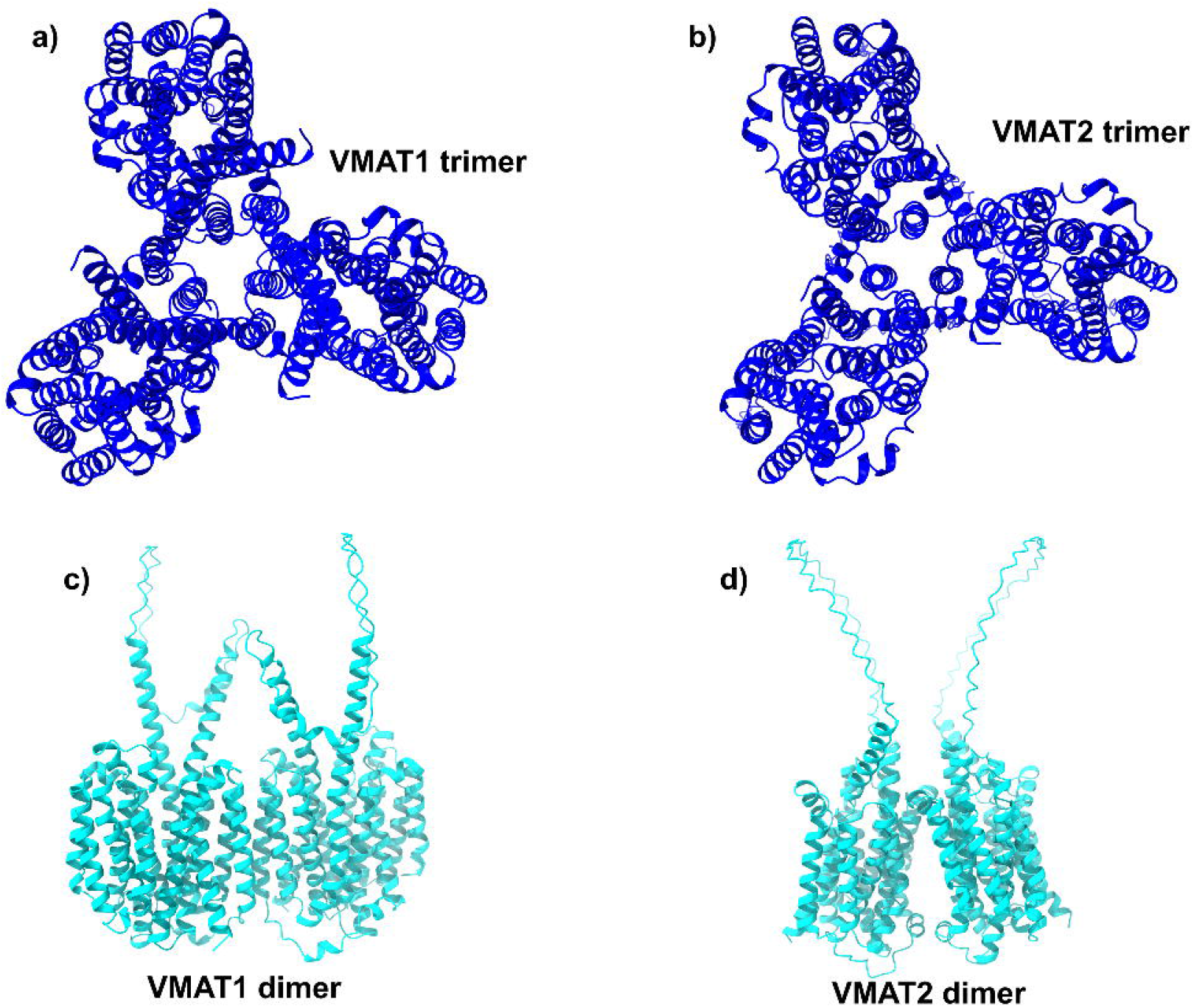
Predicted VMAT1 and VMAT2 canonical bio-assemblies. a) Structural model of the VMAT1 trimer shown in top-view orientation, revealing a three-fold symmetric organization of monomers. b) Corresponding top-view structural model of the VMAT2 trimer, exhibiting a similar overall trimeric architecture. c) Side-view representation of the VMAT1 dimer, highlighting transmembrane helix packing and interface contacts within the membrane region. d) Side-view representation of the VMAT2 dimer, illustrating comparable dimeric interactions and helical arrangements.

The truncated isoforms also exhibit distinctive biophysical features that may predict specific heterodimeric assembly mechanisms with canonical VMATs (Supplementary Table S1). Q96GL6 demonstrates elevated hydrophobicity (GRAVY 0.702) and increased basicity (pI 6.59), suggesting preferential partitioning into ordered lipid domains. This 8-TM variant retains critical binding residues within TM5-6, positioning it as a possible negative regulator that could create asymmetric functional assemblies through complementary domain interactions. The 4-TM fragment E5RK12 maintains hydrophobic character (GRAVY 0.576) despite extensive truncation, consistent with a stable membrane-inserted module capable of engaging canonical VMATs through conserved helix-helix interaction motifs.

### Comparative Docking of VMAT1 and VMAT2 Isoform Complexes

MMGBSA docking analyses (Supplementary Table S2, Supplementary Figure S7) revealed that VMAT2 binds substantially weaker to its isoform (MMGBSA ΔG ≈ -80 kcal/mol) compared to VMAT1-isoform bindings (MMGBSA ΔG ≈ -157 kcal/mol). This nearly twofold difference highlights a selective affinity of the isoforms for VMAT1, supporting the idea that truncated variants are specialized regulatory partners rather than general inhibitors of vesicular monoamine transporters. The stronger VMAT1-isoform binding may arise from isoform-specific charge complementarity and transmembrane hydrophobic continuity. This differential binding preference also aligns with the known expression divergence between the two transporters.

### Free Energy Calculations in the Synaptic Membrane Environment

Full-atom MMPBSA calculations performed over the final 50 ns of the membrane-embedded trajectories revealed distinct energetic profiles among the VMAT1 dimer, the VMAT1-isoform complex, and the isoform dimer (Fig. 2). The VMAT1 dimer exhibited the most stable total binding energy, averaging around -24.15 kcal/mol, characterized by moderate fluctuations and a consistent energetic baseline across the evaluated frames (Supplementary Figure S8,S9). This stability indicates a tightly coupled protomer interface, suggesting that the full-length VMAT1 monomers maintain a robust dimeric conformation within the lipid bilayer, consistent with the transporter’s functional requirement for synchronized conformational transitions during vesicular monoamine loading.

**Fig 2.**
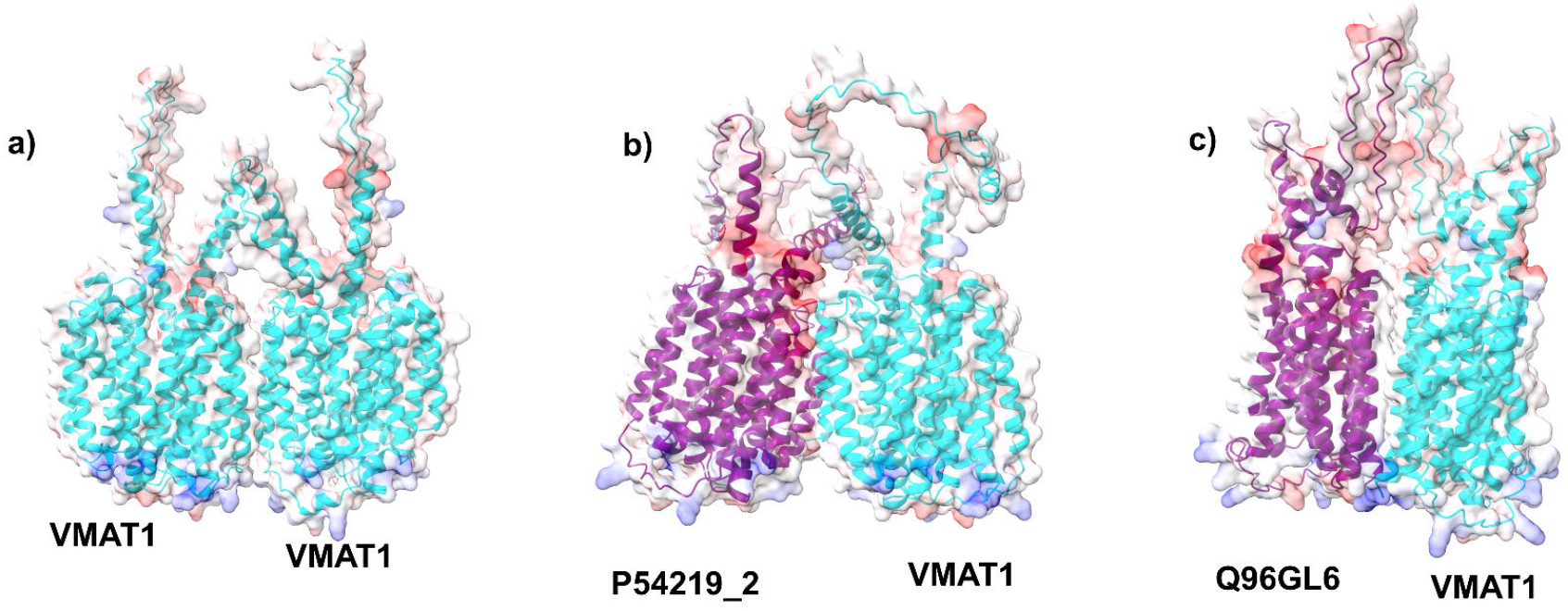
Inhibition of canonical VMAT1 dimer via truncated isoforms. a) VMAT1-VMAT1 dimer model shown with surface electrostatic projection, demonstrating the native homomeric interface and transmembrane packing stability. b) Predicted heterodimeric assembly between VMAT1 and the P54219_2, illustrating a compatible membrane-embedded interface and structural complementarity across helices.c) Docking model of VMAT1 with the VMAT-like protein Q96GL6, revealing a stable alternative heteromeric configuration with notable helix-helix interaction surfaces.

In contrast, a VMAT1-isoform complex showed a more negative and variable energetic profile, with total binding energies fluctuating between -20 to -52 kcal/mol (Fig. 2). The VMAT1-Q96GL6 heterodimer exhibited a state of thermodynamic hyper-affinity (Supplementary Figure S10,S11). This stability significantly surpasses that of the canonical VMAT1 homodimer. The resulting differential binding energy (Δ(ΔG) =-14 kcal/mol) establishes a steep thermodynamic gradient that strongly favors the formation of the heterodimer. These findings are consistent with the hypothesis that isoform incorporation into the VMAT1 oligomeric complex may act as a negative modulator, potentially disrupting vesicular packaging efficiency or trafficking dynamics in monoaminergic synapses.

### Isoform-P Homodimerization as a Protective Self-Sequestration Mechanism

The VMAT1-P54219-2 heterodimer occupies the lowest tier of the stability hierarchy, displaying a significantly attenuated mean interaction energy of -10.30 kcal/mol (Supplementary Figure S12,S13). The isoform dimer displayed the greatest energetic instability (Fig. 3), indicating a transient and energetically unfavorable dimerization pattern. This suggests that, in the absence of the full-length VMAT1 subunit, the isoform dimer cannot sustain stable membrane embedding or interfacial complementarity. Such behavior supports the view that the truncated P-isoform is not evolutionarily optimized for independent oligomerization but rather againsy competitive interaction with VMAT1, thereby modulating its assembly, localization, or functional output. Further supporting this, the isoform-P homodimer exhibits a markedly higher binding affinity, with an average interaction energy of −17.43 kcal/mol, exceeding that of the VMAT1-isoform-P heterodimer (Supplementary Figure S14,S15,S16,S17). This self-limiting behavior provides a regulatory layer: isoform-P can interact with VMAT1 when present, but its default state is to neutralize its own disruptive potential by forming homodimers. A similar mechanism is predicted for glutamate transporter isoforms [16].

**Fig 3.**
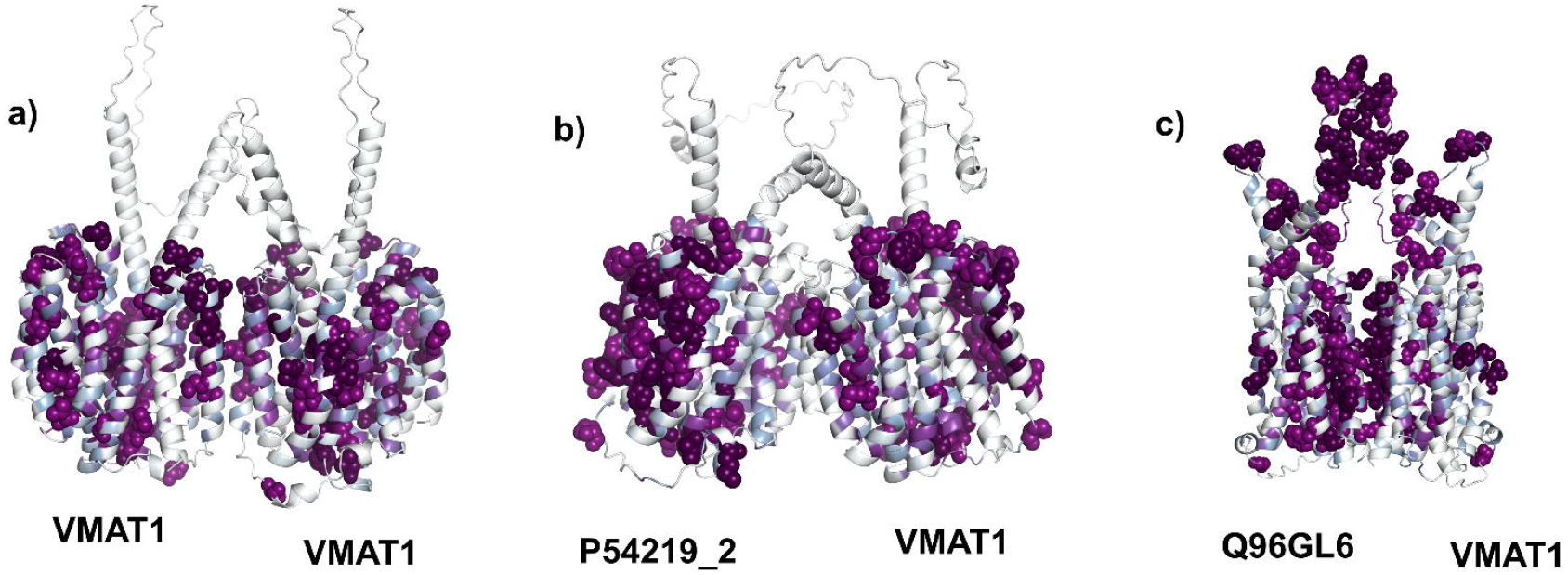
Evolutionary coupling analysis between VMAT canonical protein and splice isoforms. For Q96GL6 - Canonical VMAT1 complex, there is a strong co-evolution (colored purple) of the residues contributing to the binding interfaces.

### Mechanisms of Specificity: The Electrostatic Latch and Hydrophobic Packing

To resolve the atomic-level drivers of this energetic divergence, Per-Residue Energy Decomposition (PRED) was employed. This analysis revealed a fundamental mechanistic switch between the non-specific hydrophobic packing of the canonical form and the precise electrostatic steering of the inhibitory Q96GL6-isoform. The hyper-stability of the VMAT1-Q96GL6 heterodimer is driven by an electrostatic interaction network that functions as a molecular latch. The decomposition heatmaps highlighted a high-intensity interaction cluster anchored by residues B:Glu209 and Arg211. Glu209, a residue unique to the Q-isoform interface contribution, acts as a primary acidic anchor, exhibiting massive favorable energy contributions (-40 kcal/mol). This interaction is further stabilized by Asp205, creating an anionic pocket of high charge density. The enthalpic gain from this specific coulombic attraction is sufficient to override local steric frustrations and repulsive penalties (e.g., Arg212), locking the complex into a rigid, low-energy conformation.

Conversely, the canonical VMAT1 homodimer relies on a distributed network of hydrophobic contacts typical of transmembrane helix interactions. The stabilizing forces are mediated primarily by residues such as B:Leu245, A:Phe170, and B:Met167 (A:VMAT1, B:Ligand). The binding energy in this system is driven largely by the entropic gain associated with solvent exclusion and the favorable enthalpy of van der Waals packing. The instability of the VMAT1-P heterodimer can be attributed to a failure of interfacial complementarity. Isoform P lacks the acidic Glu209 anchor required for electrostatic locking, yet also fails to present a coherent hydrophobic surface capable of efficient packing against VMAT1. The decomposition profile reveals a diffuse, low-energy landscape with significant contributions from destabilizing interactions, such as steric clashes involving A:Asn436.

### David vs. Goliath

This mechanism “solves” the David vs. Goliath conflict by changing the rules of engagement from a battle of size (surface area) to a battle of intensity (charge). In a standard biological context, the larger protein (“Goliath”) would naturally dominate because binding usually scales with surface area: the more contact points, the more stable the complex. A smaller fragment (“David”) inherently lacks the structural mass to compete on these terms and would typically be crowded out. Instead of trying to match the canonical protein’s broad hydrophobic surface, the isoform concentrates its binding energy into high-potency “electrostatic latch” (Glu209-Arg211). By achieving a binding energy of -37 kcal/mol (vs. the canonical -23 kcal/mol), the isoform creates a “thermodynamic trap”. This solves the problem of competition; the system is energetically forced to favor the heterodimer, guaranteeing that the “David” wins every time it encounters a “Goliath”, effectively functioning as a negative regulator.

### Co-evolutionary Profilling of VMAT Isoform Regulation

Molecular evolution of transmembrane proteins is non-linear and complex [24,26] and involves co-evolutionary dependencies [26,27,28]. The existence and structural specificity of truncated VMAT1 isoforms suggest that these variants are not merely transcriptional errors or targets for nonsense-mediated decay, but rather evolved regulatory modules that function as “splicing-sensitive checkpoints” in monoaminergic signaling. This is structurally evidenced by the mechanism observed in the Q-isoform (Q96GL6), which utilizes a precise high-affinity interaction of Glu209 and Arg211 to override the canonical hydrophobic dimerization interface. Based on the evolutionary couplings and the specific biophysical properties of the Q-isoform, this structural mapping delineates a highly integrated epistatic network of residues undergoing correlated mutations to maintain a specific functional architecture under stringent purifying selection.

The dense clustering of co-evolving residues along the transmembrane interface signifies that the “electrostatic latch” mechanism is not a stochastic occurrence but a conserved interaction motif (Fig. 3). Furthermore, the extension of this co-evolutionary network into the extracellular (luminal) domains suggests that the inhibitory interface is allosterically coupled to the transporter’s gating machinery or extracellular signalling activity. This implies that the extracellular loops condition the binding interface, ensuring that the dominant-negative sequestration is conformationally selective, likely locking the transporter in a specific state to enforce the regulatory checkpoint effectively within the complex synaptic vesicle membrane environment. This may also suggest a potential signalling function. This indicates that the potential for isoform-mediated inhibition is a selected trait in VMAT1, likely evolved to fine-tune serotonergic and noradrenergic tone, whereas VMAT2 has been selected for stability and resistance to such modulation to preserve the integrity of critical neurotransmission.

### Structural Implications for Therapeutic Targeting

The energetic destabilization of the VMAT1-isoform complex in membrane MMPBSA simulations, coupled with its high binding affinity in docking analyses, implies a dual regulatory mechanism: the isoform strongly associates with VMAT1 but induces local conformational stain. Such negative modulation could contribute to reduced monoamine storage. In contrast, the relatively weak isoform-VMAT2 interaction predictions indicates that VMAT2 is functionally insulated from isoform interference, preserving dopaminergic signaling integrity. This molecular selectivity has drug-targeting implications, as pharmacological agents that stabilize VMAT1 dimer interfaces or prevent isoform binding.

The unique Glu209/Arg211 electrostatic axis of the inhibitory VMAT1-Q isoform complex presents a high-specificity pharmacophore for rational drug design, distinct from the hydrophobic surface of the canonical transporter. The most direct therapeutic strategy involves the design of “decoy” inhibitors to rescue canonical VMAT1 function. Hydrocarbon-stapled alpha-helical peptides, engineered to mimic the VMAT1 interfacial helix containing Arg211, could competitively bind to the acidic Glu209 pocket of Isoform Q. By capping this electrostatic site, such peptidomimetics would prevent the sequestration of VMAT1 monomers, shifting the thermodynamic equilibrium back toward the formation of functional homodimers.

The specific anionic cluster (Asp205/Glu209) on Isoform Q offers an ideal recognition motif for PROTAC development. Alternatively, the canonical VMAT1 homodimer itself can be targeted via “molecular glues” designed to bind the hydrophobic cleft defined by Leu245 and Phe170. A bivalent stabilizer binding at the dimer interface could contribute additional binding enthalpy, lowering the binding energy of the homodimer to levels competitive with the Q-heterodimer. This allosteric reinforcement would render the functional assembly energetically robust against competitive inhibition by Isoform Q.

### Conclusion

Together, these results reveal that the isoform preferentially binds to VMAT1, but not VMAT2, within the synaptic membrane environment. This selective interaction suggests that VMAT1 is uniquely susceptible to isoform-mediated conformational modulation, providing a potential mechanistic explanation for the transporter’s tissue-specific regulation and vulnerability in mood disorders. The study further posits that the Q-isoform has specifically evolved to inhibit the VMAT1 assembly process through this high-fidelity electrostatic steering, acting as a negative modulator or signalling complex that creates a state of thermodynamic hyper-affinity compared to the homodimer. This implies that alternative splicing in VMAT1 has been co-opted as a post-transcriptional strategy to dynamically limit vesicle storage capacity during states of synaptic plasticity or metabolic stress, providing a rapid mechanism to downregulate neurochemical signaling without altering genomic expression. The findings highlight the isoform-VMAT1 axis as a potential therapeutic target for restoring vesicular monoamine homeostasis through selective stabilization of VMAT1 oligomeric architecture.

## Methods

### Protein Sequences and Other Characteristics

Protein sequence data was acquired in FASTA format from UniProt [12]. Corresponding protein-coding isoforms were identified using the Uniprot, which employs automatic gene-centric mapping derived from eukaryotic reference proteomes and original sequencing projects [12]. This analysis specifically targeted non-decayed protein-coding isoforms with lengths ranging from 15% to 85% of the canonical full-length sequence. Membrane topology and sequence features were visualized via the Protter web application [29]. Physicochemical properties, including molecular weight (MW), amino acid composition, instability indexes, and GRAVY scores, were calculated using Expasy [30,31,32]. The AlphaFold3 server was employed to predict monomeric, dimeric, and trimeric structures [33].

### AlphaFold3 predictions and model ranking

Given the study’s emphasis on the accurate ranking of inhibitors, the computationally efficient Molecular Mechanics with Generalized Born and Surface Area (MM/GBSA) method was selected [34]. Complex re-ranking was performed using the HawkDock web server [34,35,36,37], following the addition of missing hydrogens and heavy atoms via the tleap module in Amber16. Calculations relied on variable dielectric generalized Born model, which incorporates residue-type-based dielectric constants and outperforms classical generalized Born models [35]. The system underwent a 5,000-step minimization (2,000 steps of steepest descent followed by 3,000 steps of conjugate gradient) with a 12 Å cutoff for van der Waals interactions. Finally, free energy decomposition was utilized to highlight key residues in the receptor (canonical transporter) and ligand (isoform) interface.

### Atomistic synaptic vesicle membrane systems

According to the methodology explained in our previous studies [16,25,38,39], all-atom membrane systems were constructed using the CHARMM-GUI Membrane Builder [40,41,42]. Protein spatial alignment within the lipid bilayers was optimized via the PPM 2.0 algorithm, which incorporates hydrogen bonding and anisotropic dielectric properties at the membrane-water interface [43]. The lipid composition was designed to mimic a representative plasma membrane, consistent with our previous models [16,17,21,22]. The membrane contained cholesterol, POPC, POPE, POPS, POPI, and palmitoyl-sphingomyelin (PSM) lipids distributed symmetrically across both leaflets [21,22]. System neutrality was established by adding K□ and Cl□ ions to an ionic strength of 0.15 M, determined through 2,000 steps of Monte Carlo simulation using a primitive ion model. The CHARMM36m all-atom force field was applied throughout [44].

### Molecular dynamics simulations

Molecular dynamics simulations were executed on AlphaFold-predicted membrane structures using GROMACS 2024.3 [45], adhering to protocols established in our prior studies on glutamate transporter variants and neuronal proteins [16,25,387]. Computational analyses utilized a cluster of three Google Colab instances, equipped with an NVIDIA L4 GPU, 126 GB of VRAM, 318 GB of RAM, and Intel® Xeon® CPUs [46]. To maximize efficiency, the software was recompiled with CUDA support, enabling parallelization across multiple cores [46]. Energy minimization was performed using the steepest descent algorithm for a maximum of 5,000 steps. Convergence was defined by a maximum force tolerance of 1000 kJ/mol/nm. Equilibration followed the standard six-step CHARMM-GUI protocol, characterized by the gradual release of restraints. The final equilibration step consisted of a 500 ps run utilizing the v-rescale thermostat to maintain a temperature of 303.15K (τ t =1.0ps) and the C-rescale barostat to maintain a pressure of 1bar via semi-isotropic coupling (τp=5.0ps). Production MD simulations were carried out for 50 ns with frames saved every 0.5 ns, consistent with our earlier studies [16]. Electrostatic interactions were computed via the Particle Mesh Ewald (PME) method, applying a 1.2 nm cutoff for both Coulomb and van der Waals terms. The system temperature (303.15 K) and pressure (1 bar) were maintained using the Nose-Hoover thermostat and the Parrinello-Rahman barostat under semi-isotropic coupling. Hydrogen bonds were constrained using the LINCS algorithm, allowing for a 1.2 nm cutoff for short-range van der Waals and electrostatic interactions.

System stability was evaluated through trajectory analysis. The root mean square deviation (RMSD) of the protein backbone and the radius of gyration (Rg) were calculated using the GROMACS rms and gyrate modules, respectively. Residue flexibility was assessed by computing the root mean square fluctuation (RMSF) of C-alpha atoms. Solvent accessible surface area (SASA) for protein side chains was determined using the gmx sasa tool with a probe radius of 1.4 Å [47]. For binding free energy estimations, the final 20 ns (31-50 ns) of the equilibrated trajectories were analyzed.

### Binding Free Energy Calculations

Binding free energies were computed using the Molecular Mechanics Poisson-Boltzmann Surface Area (MMPBSA) method via gmx_MMPBSA for membrane systems [48,49]. Dielectric constants were assigned as follows: membrane (7.0), solute (4.0), and solvent (80.0). Solvent-excluded surfaces were defined using a 1.40 Å water probe and a 2.70 Å membrane probe. Total electrostatic energies and forces were evaluated using the particle-particle particle-mesh (P3M) method. Per-residue energy decomposition (idecomp = 2) was applied to distinguish individual contributions to electrostatic and van der Waals components. Binding energies were averaged over the selected time steps, with the standard error of the mean (SEM) calculated via the propagation of uncertainty. Interface contacts (<4 Å) were analyzed and visualized using gmx_MMPBSA_ana and UCSF ChimeraX [50], while plots were generated using Grace (https://plasma-gate.weizmann.ac.il/Grace/).

### Co-evolutionary Profilling

Evolutionary couplings (ECs) were calculated via the EVcouplings server (http://evfold.org) using a maximum entropy model [16,51] constrained by multiple sequence alignment (MSA) statistics. To allow for robust comparisons across varying protein lengths and database sizes, length-normalized bitscores were employed. The quality of the alignment was assessed using the ratio of effective sequences to protein length (n_eff/L), where a value greater than 5 generally indicates a high-quality run. The calculated n_eff /L ratios were 86.68 for canonical VMAT1, 123.23 for the P-isoform (P54219-2), 14.16 for the Q-isoform (Q96GL6), and 2.30 for the E5RK12 isoform.

## Supporting information

Sumpplementary File

## Ethics Approval

Ethics approval was not required for this computational study as it did not involve animal subjects, human participants, and identifiable data.

## Consent to participate

Not applicable. This computational study did not involve human participants.

## Consent for publication

Not applicable. This computational study did not involve human participants.

## Availability of data and materials

The AlphaFold DB (https://alphafold.ebi.ac.uk), a database developed by DeepMind and the European Bioinformatics Institute (EMBL-EBI) at EMBL, is a repository for AlphaFold2 predictions, with over 200 million protein structures. Each statistical and computational analysis of this study, included with step-by-step instructions where possible, are publicly available to ensure repeatability. For more detailed information on the statistical analyses, input files and detailed outputs, including the AlphaFold calculations and codes to regenerate analyses, please visit the website:https://github.com/karagol-alper/VMAT-isoforms

## Author Contributions

A.K and T.K contributed equally to this work. All authors have read and agreed to the published version of the manuscript.

## Acknowledgements

None.

---

**Table 1.** **Interface composition and binding free energies (**ΔΔ**Gs) of the dimer complexes in an atomistic bilayer model**.

| Receptor | Ligand<sup>1</sup> | Average Binding free energy ($\Delta\Delta G$) (kcal/mol)<sup>3</sup> | SD (standard deviation) | Thermodynamic Assessment vs. Canonical |
| --- | --- | --- | --- | --- |
| **VMAT1** | **VMAT1 (DIMER)** | -24.15 | 4.91 | Baseline Stability |
| | **P54219_2** | -10.30 | 5.20 | $\Delta\Delta G=+13.85$ kcal/mol (Highly Disfavored) |
| | **Q96GL6** | -37.36 | 6.93 | $\Delta\Delta G=-14.21$ kcal/mol (Highly Favored) |
| **P54219_2** | **P54219_2 (DIMER)** | -17.43 | 3.63 | Less Stable than Canonical |
| **Q96GL6** | **Q96GL6 (DIMER)** | -25.48 | 8.90 | Slightly More Stable than Canonical |

<sup>1</sup>The Uniprot Entry ID of the isoform (ligand of the complex).
<sup>2</sup>Residue contribution of the binding energy, sorted by calculated energies (kcal/mol), the highest contributing (lowest energy) first 5 residues listed.
<sup>3</sup>Residue binding free energies are calculated according to MMPBSA algorithm.

