## Supplementary material for "Splice Isoform Induced Selective Inhibition of Vesicular Monoamine Transporter Assembly Revealed by Multi-level Combinatorial Analysis": Sumpplementary File

**Table S1. Computationally mapped and predicted potential truncated isoforms of vesicular monoamine transporter subfamily (VMATs).**

| **Name** | **Isoform ID^1^** | **TM count^2^** | **Length (aa)^3^** | **Mass (Da)^4^** | **Isoelectric point (pI)** | **Instability index^5^** | **GRAVY^6^**  **^(cut-off=0)^** |
| --- | --- | --- | --- | --- | --- | --- | --- |
| **VMAT1** | [P54219](https://www.uniprot.org/uniprotkb/P54219/entry) (Canonical) | 12 | 525 | 565257 | 5.59 | 44.45 | 0.585 |
|  | [P54219-2](https://www.uniprot.org/uniprotkb/P54219/entry) | 11 | 472 | 50026 | 5.58 | 44.35 | 0.583 |
|  | [Q96GL6](https://www.uniprot.org/uniprotkb/Q96GL6/entry) | 8 | 385 | 41172 | 6.59 | 40.79 | 0.702 |
|  | [E5RK12](https://www.uniprot.org/uniprotkb/E5RK12/entry) | 4 | 258 | 27669 | 5.64 | 42.17 | 0.576 |
| **VMAT2** | [Q05940](https://www.uniprot.org/uniprotkb/Q05940/entry) (Canonical) | 12 | 514 | 55713 | 5.68 | 48.36 | 0.579 |
|  | [Q05940-2](https://www.uniprot.org/uniprotkb/Q05940/entry#Q05940-2) | 2 | 209 | 22980 | 6.89 | 47.98 | 0.383 |

^1^The Uniprot entry ID of the isoform (only included isoforms that have length between 10% and 90% of the canonical sequence). Please see Methods.

^2^Transmembrane (TM) domain count of the protein, derived from the topology information included in the Uniprot entries.

^3^The amino acid length of the isoform sequence.

^4^Protein mass calculated from sequences.

^5^The instability index calculated from input sequence as an estimate of the stability of the protein in a test tube, according to correlation analysis of Guruprasad et. Al (Methods).

^6^The GRAVY (Grand Average of Hydropathy) value for the corresponding protein is calculated as the sum of hydropathy values of all the amino acids, divided by the number of residues in the sequence.

**Table S2. Interface composition and binding free energies (ΔΔGs) of the sampled dimer complexes.**

| **Receptor** | **Ligand^1^** | **Contact Composition^2^** | | **Total binding free energy (ΔΔG) (kcal/mol)^3^** |
| --- | --- | --- | --- | --- |
|  |  | **Receptor** | **Ligand** |  |
| **VMAT1** | **VMAT1 (DIMER)** | PHE-170, LEU-259, VAL-166, LEU-252, LEU-245 | PHE-170, LEU-259, VAL-166, LEU-252,VAL-159 | -145.03 |
|  | **P54219_2** | CYS-459, ASN-452, PHE-441, MET-445, TYR-455 | GLU-431, LEU-365, LEU-364, TRP-310, MET-357 | -254.56 |
|  | **Q96GL6** | ARG-211, TRP-320, GLN-5, PHE-374, HIS-207 | ARG-212, GLN-6, TRP-321, LYS-347, HIS-208 | -290.03 |
| **VMAT2** | **VMAT2 (DIMER)** | ALA-431, TYR-167, LYS-275, THR-271, LYS-430 | ALA-431, TYR-167, LYS-275, THR-271, LYS-430 | -148.13 |
|  | **VMAT1** | PHE-253, LEU-245, LEU-238, THR-234, VAL-231 | PHE-249, LEU-252, LEU-245, LEU-259, PHE-170 | -115.97 |
|  | **P54219_2** | LEU-238, LEU-245, PHE-253, VAL-231, ALA-242 | PHE-249, LEU-245, LEU-252, VAL-238, LEU-259 | -133.76 |
|  | **Q96GL6** | PHE-9, LEU-243, PHE-253, VAL-231, LEU-246 | PHE-250, VAL-239, LEU-254, CYS-261, LEU-251 | -136.44 |
| **P54219_2** | **P54219_2 (DIMER)** | LEU-358, LEU-354, TRP-350, GLU-431, PHE-295 | LEU-354, TRP-350, LEU-358, GLU-431, PHE-295 | -159.49 |
| **Q96GL6** | **Q96GL6 (DIMER)** | TRP-321, GLN-6, ARG-212, PHE-238, LEU-325 | TRP-321, ARG-212, GLN-6, ARG-2, PHE-238, LEU-325 | -254.81 |

^1^The Uniprot Entry ID of the isoform (ligand of the complex).

^2^Resdiue contribution of the binding energy, sorted by calculated energies (kcal/mol), the highest contributing (lowest energy) first 5 residues listed.

^3^Residue binding free energies are calculated according to MMGBSA algorithm.

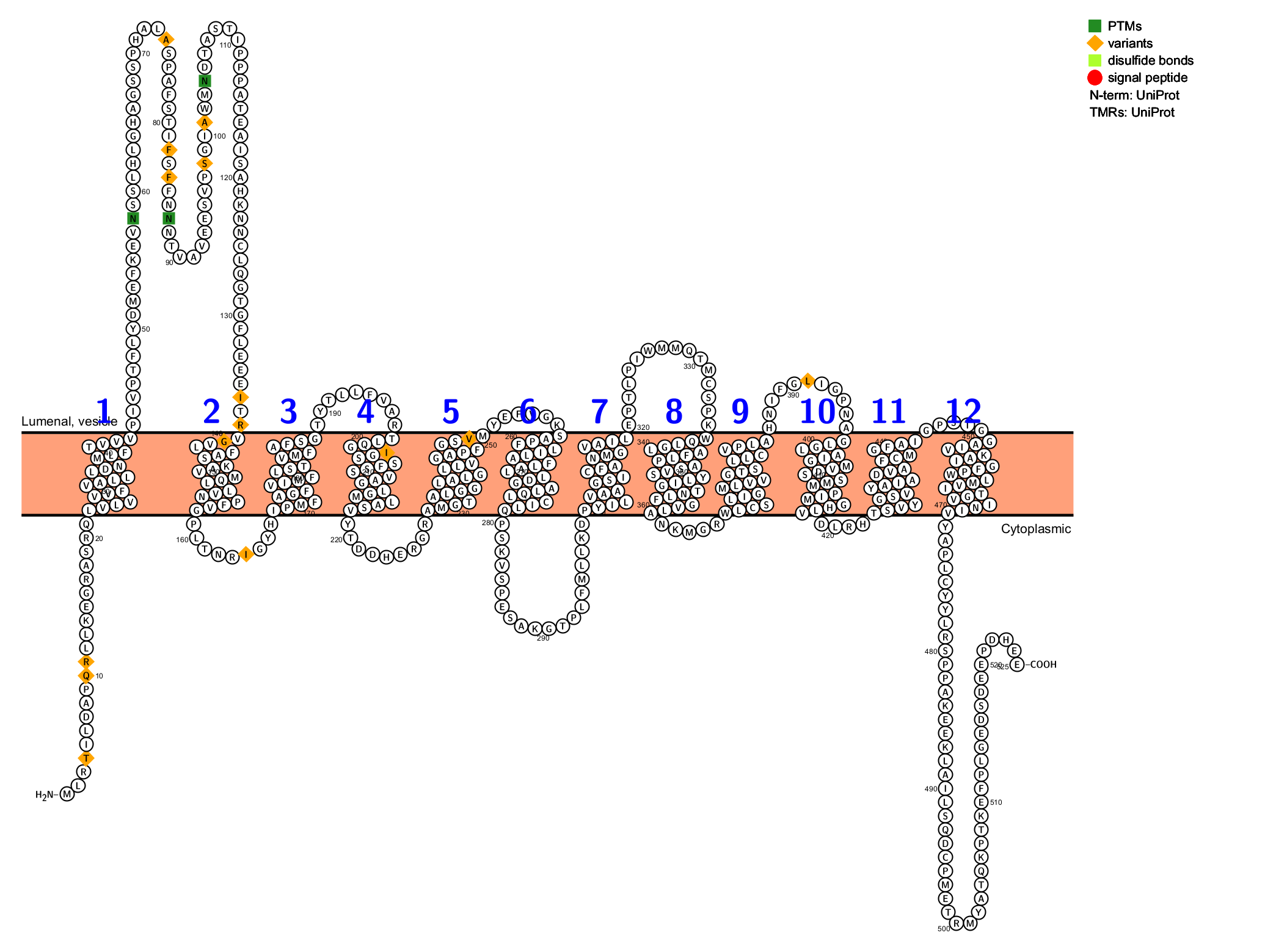
**Figure S1. Molecular topology of VMAT1 canonical isoform.** The structural data is retrieved from UniProt and visualized with Protter. For structural bioinformatic studies of VMAT1 canonical structure please also see <https://doi.org/10.1371/journal.pone.0300340>

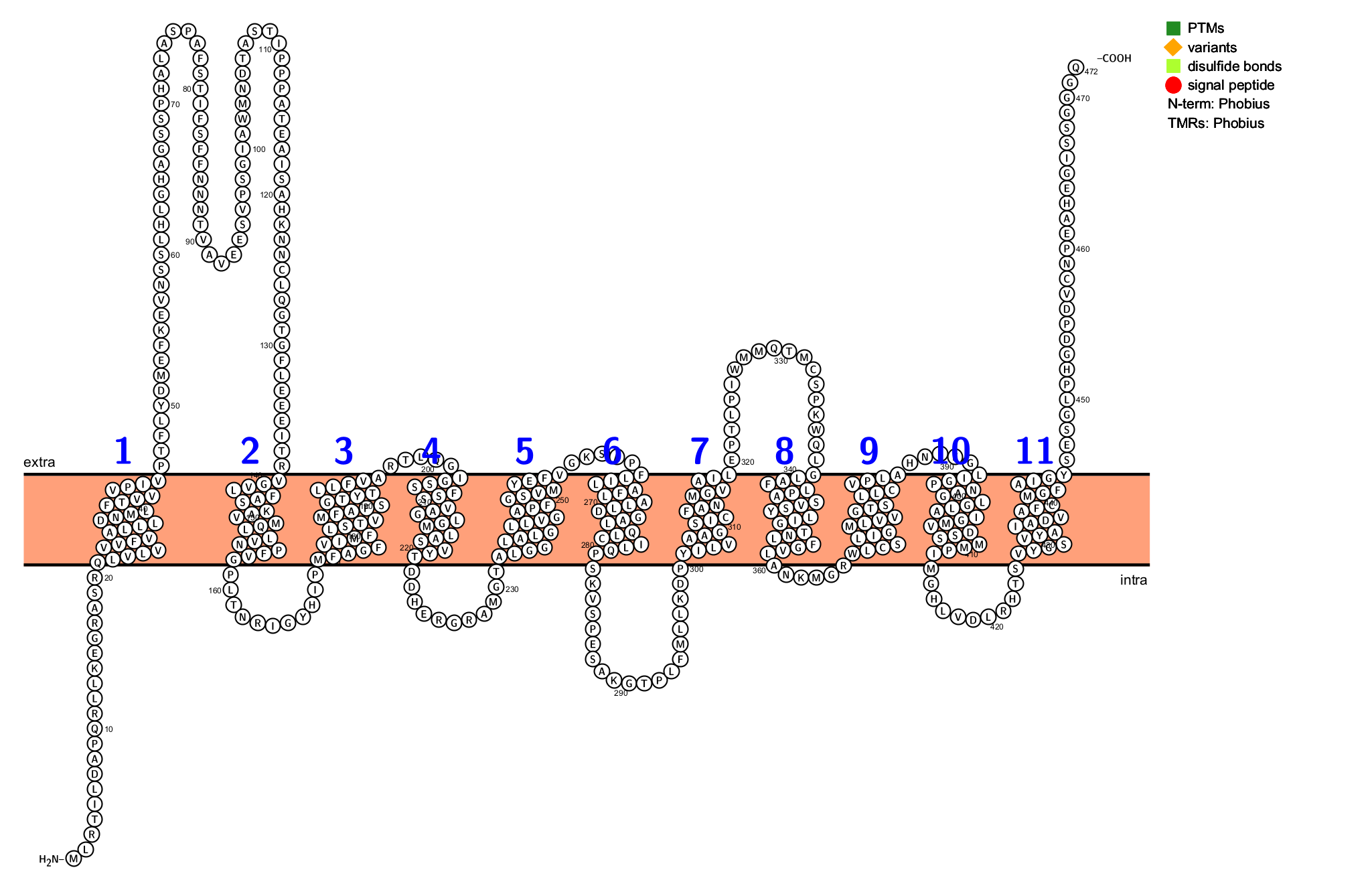
**Figure S2. Molecular topology of VMAT1 isoform P54219-2.** The structural data is retrieved from UniProt and visualized with Protter.

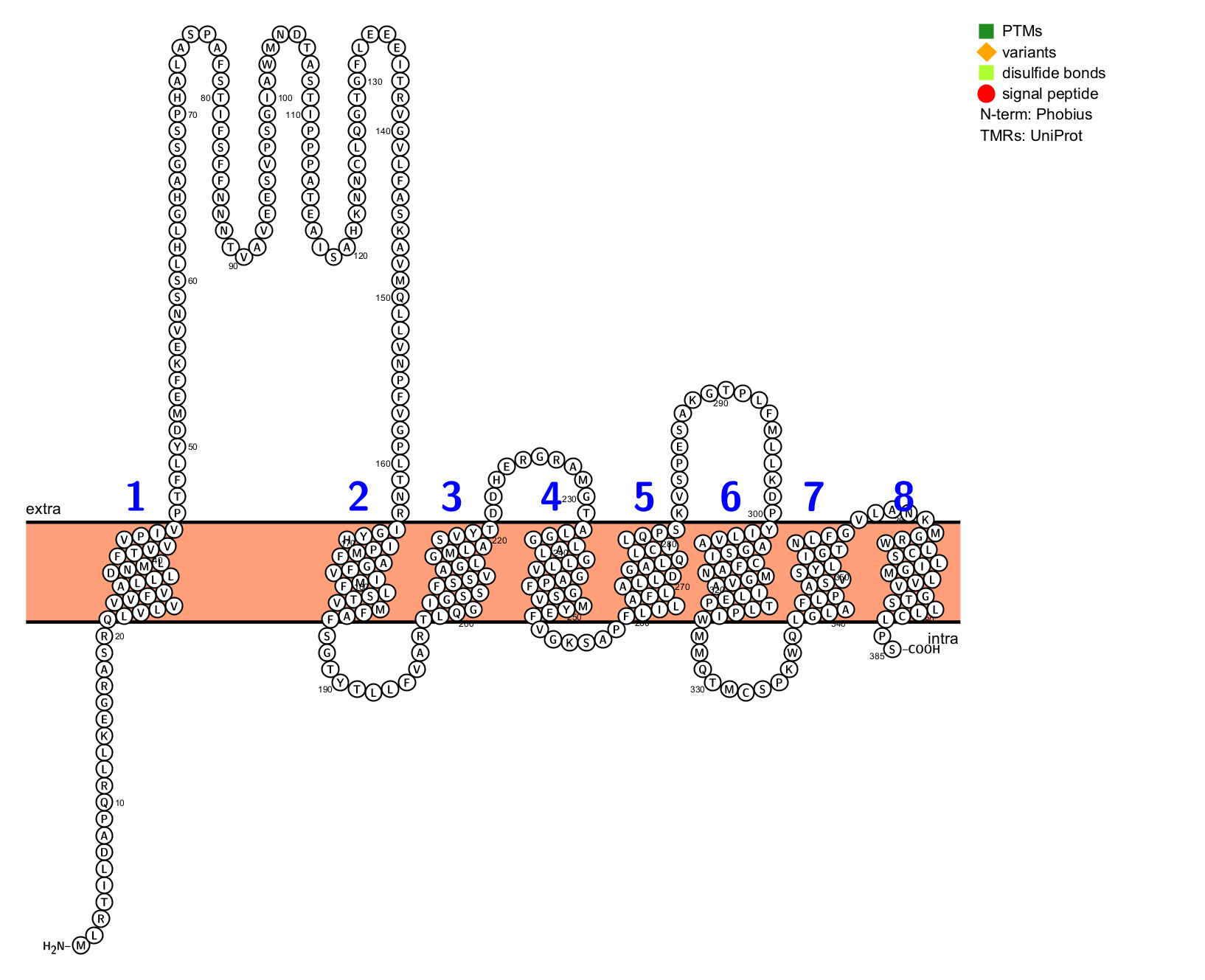
**Figure S3. Molecular topology of VMAT1 isoform Q96GL6.** The structural data is retrieved from UniProt and visualized with Protter.

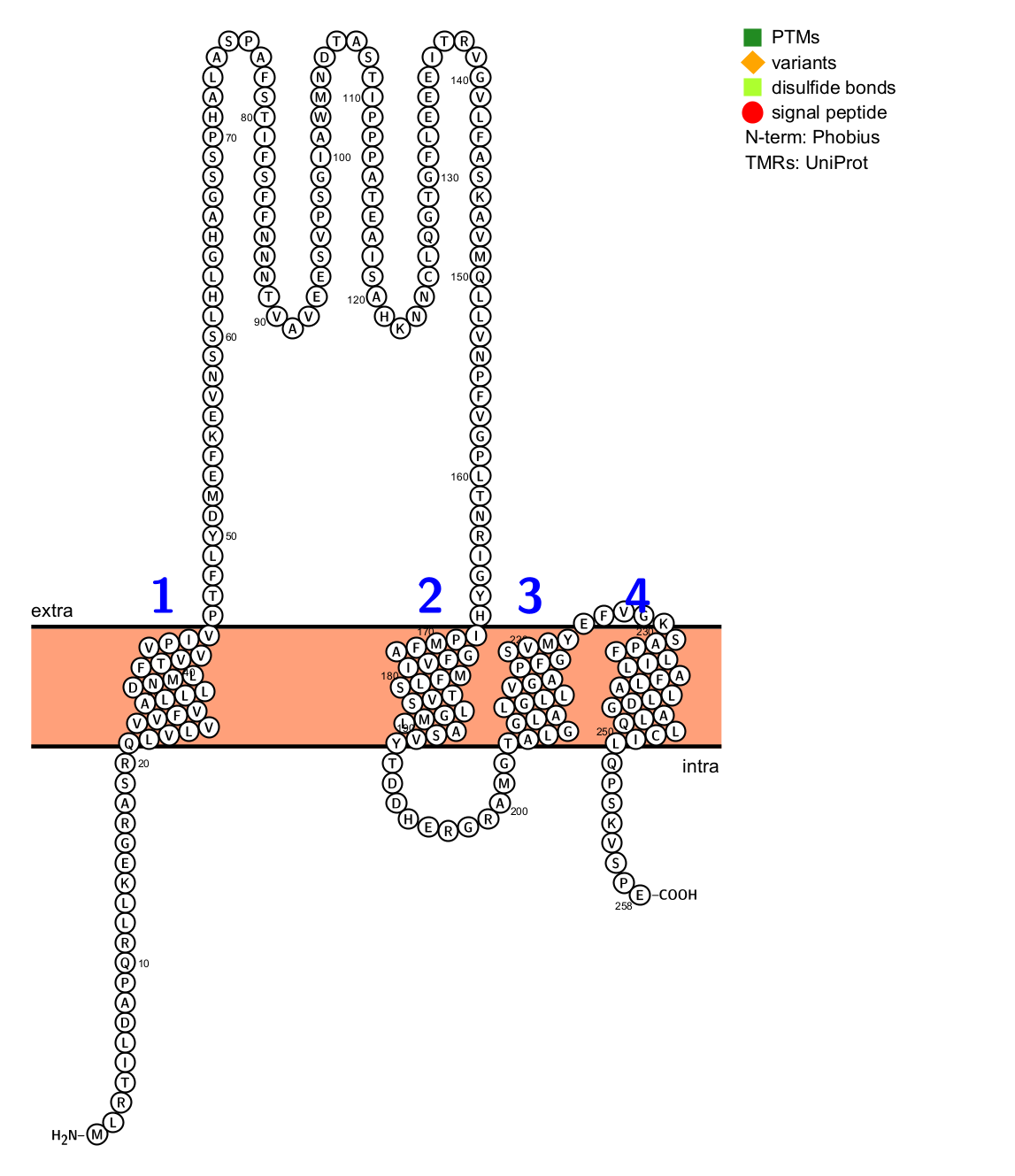

**Figure S4. Molecular topology of VMAT1 isoform E5RK12.** The structural data is retrieved from UniProt and visualized with Protter.

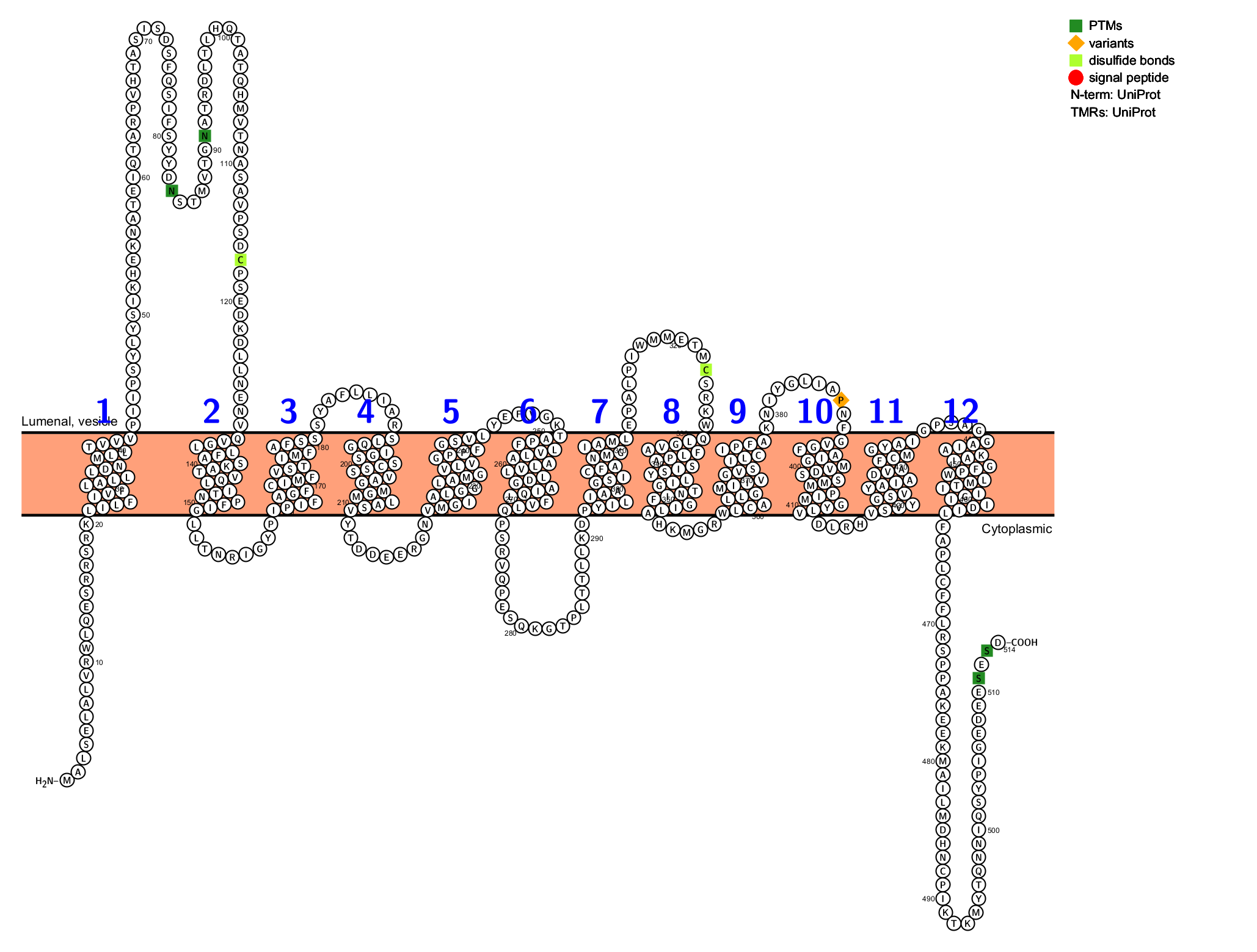

**Figure S5. Molecular topology of VMAT2 canonical isoform.** The structural data is retrieved from UniProt and visualized with Protter. For structural bioinformatic studies of VMAT1 canonical structure please also see <https://doi.org/10.1371/journal.pone.0300340>

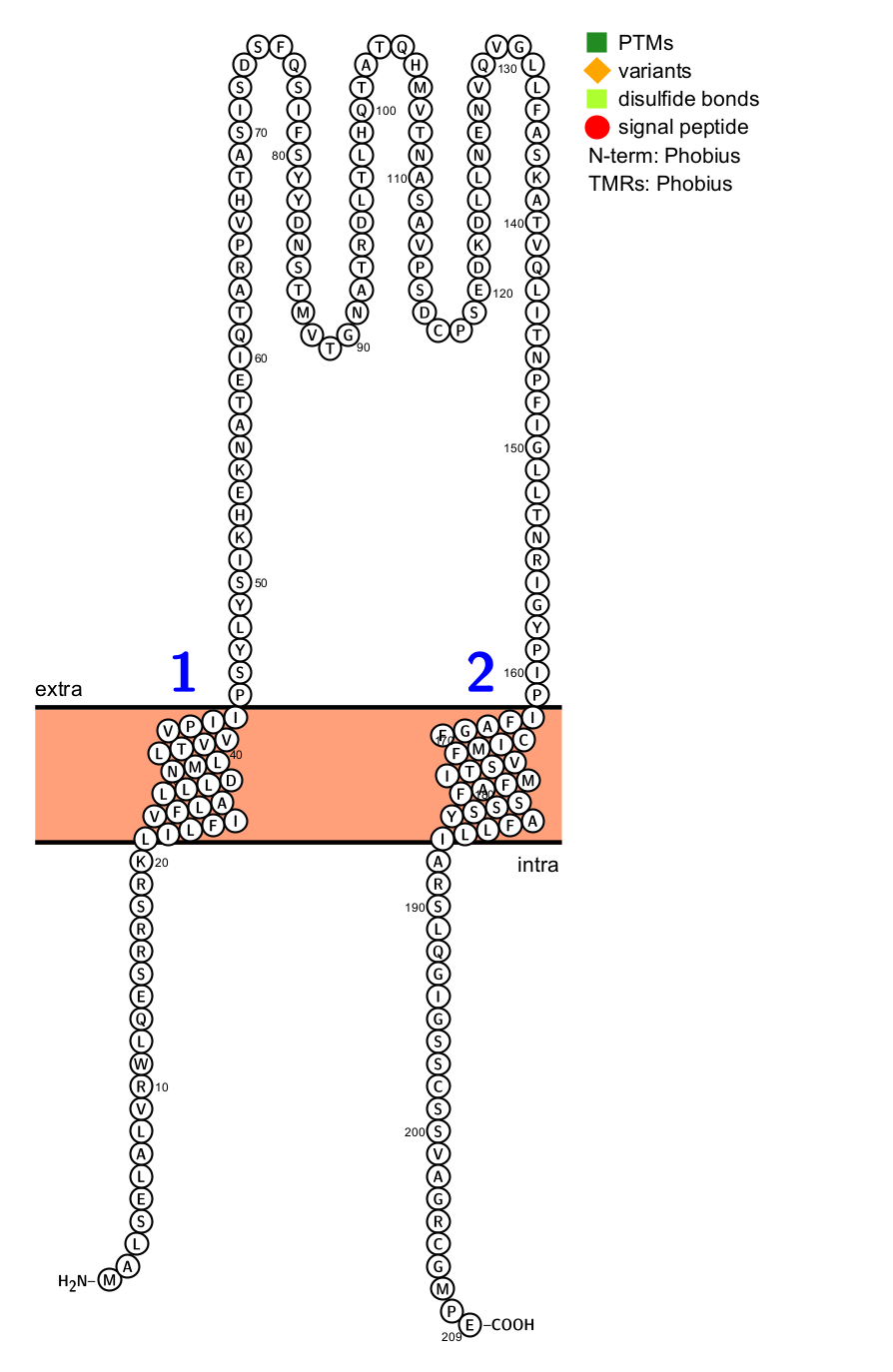

**Figure S6. Molecular topology of VMAT2 isoform Q05940-2.** The structural data is retrieved from UniProt and visualized with Protter.

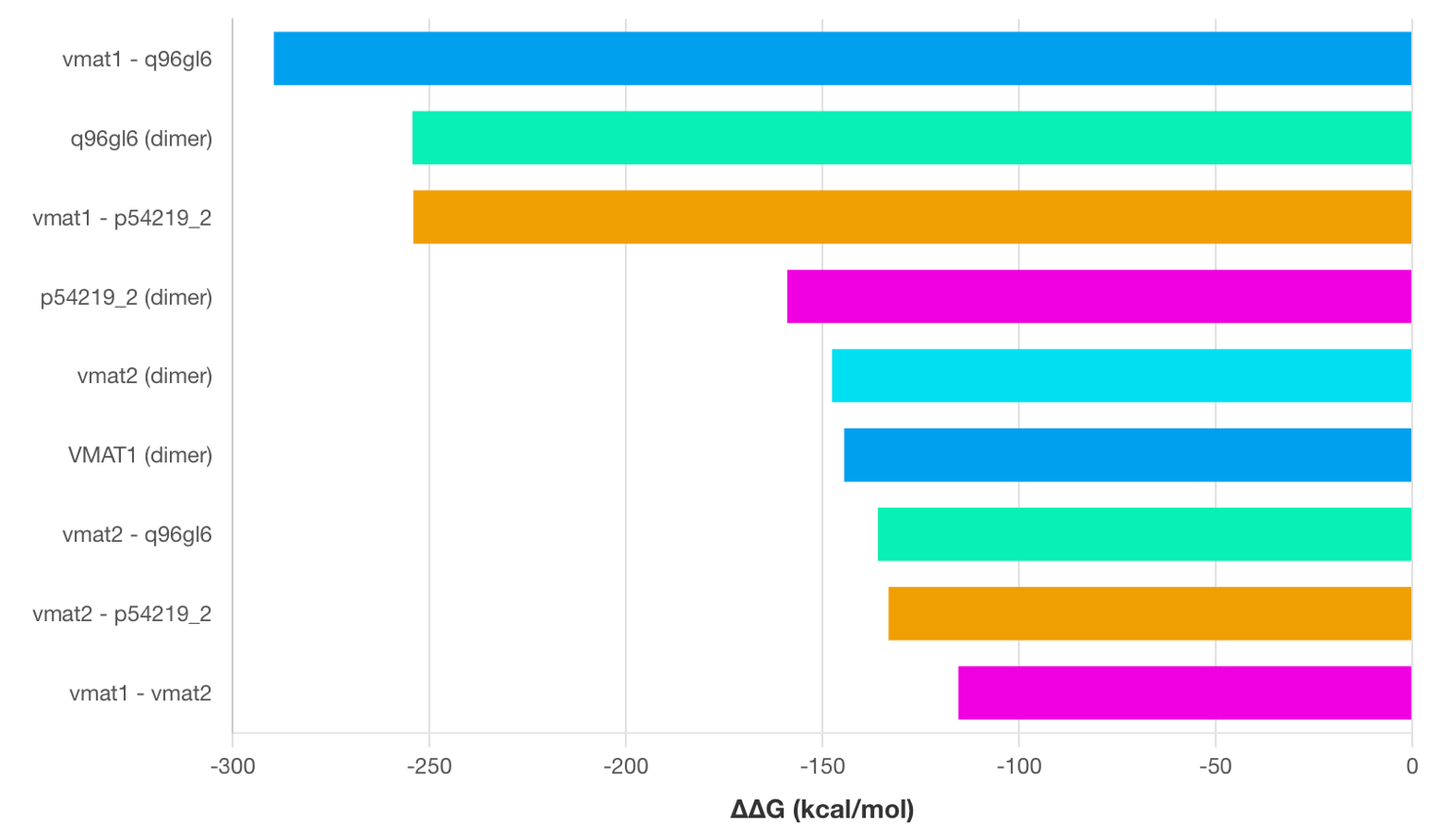

**Figure S7. MMGBSA binding free energy predictions of sampled dimers.**

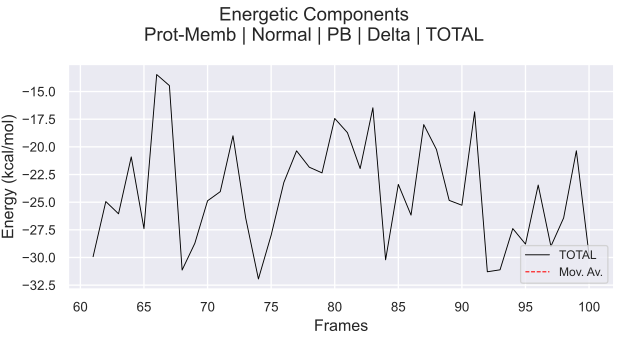

**Figure S8. Total Binding Energy Fluctuations of the VMAT1 Homodimer Across 50ns full-atom MD simulation in realistic bilayer.** The total MM/PBSA binding energy of the VMAT1 dimer calculated over simulation frames 60-100 (30-50ns). The black line represents the TOTAL binding free energy (kcal/mol), capturing frame-to-frame fluctuations in the overall interaction strength between monomers. The red dashed curve shows a moving average, highlighting underlying energetic trends and smoothing short-timescale variability. Energies include contributions from protein–membrane interactions, van der Waals forces, electrostatics, polar solvation (PB), and nonpolar terms.

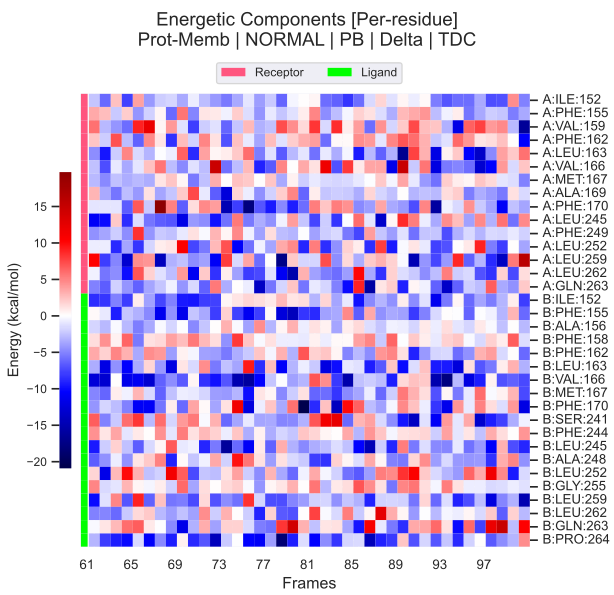

**Figure S9. Per-Residue Binding Energy Decomposition of the VMAT1 Homodimer across 50ns full-atom MD simulation in realistic bilayer.** Heatmap visualization of residue-wise MM/PBSA energy components (kcal/mol) across simulation frames 60-100 (30-50ns). Each column corresponds to a frame, and each row to an interface residue from monomer A or B. Red shades indicate energetically unfavorable contributions, while blue shades indicate stabilizing interactions. The pink and green sidebars designate receptor and ligand monomer residues, respectively (chain A vs chain B).

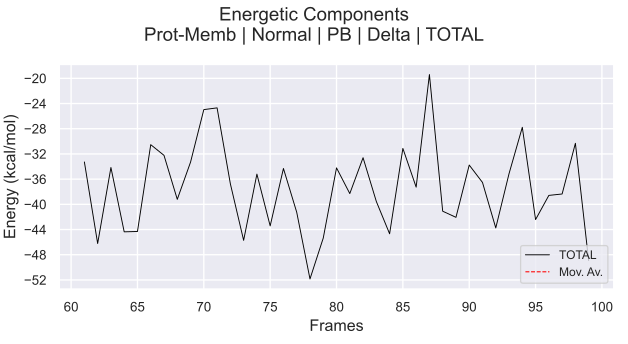

**Figure S10. Total Binding Energy Fluctuations of the VMAT1 - Isoform Q Heterodimer Across 50ns full-atom MD simulation in realistic bilayer.** The total MM/PBSA binding energy of the VMAT1 dimer calculated over simulation frames 60-100 (30-50ns). The black line represents the TOTAL binding free energy (kcal/mol), capturing frame-to-frame fluctuations in the overall interaction strength between monomers. The red dashed curve shows a moving average, highlighting underlying energetic trends and smoothing short-timescale variability. Energies include contributions from protein–membrane interactions, van der Waals forces, electrostatics, polar solvation (PB), and nonpolar terms.

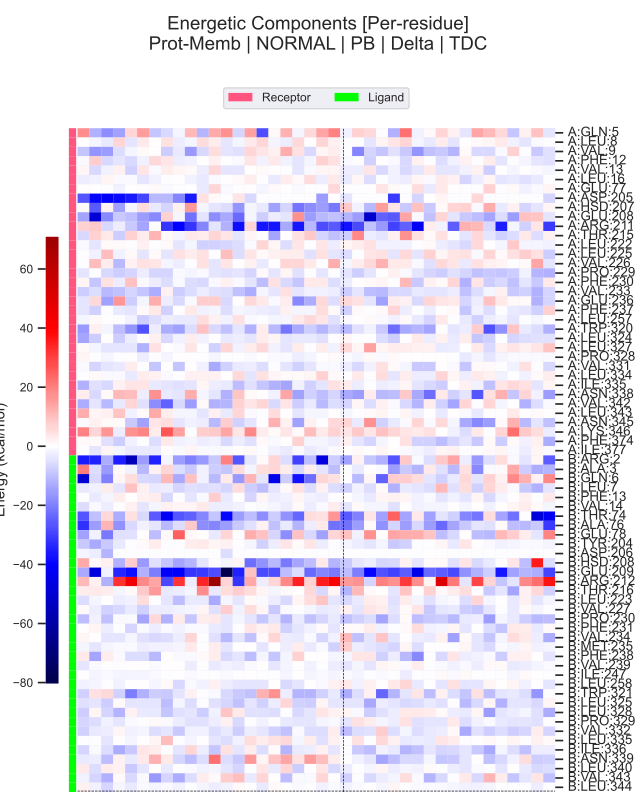

**Figure S11. Per-Residue Binding Energy Decomposition of the VMAT1 - Isoform Q Heterodimer across 50ns full-atom MD simulation in realistic bilayer.** Heatmap visualization of residue-wise MM/PBSA energy components (kcal/mol) across simulation frames 60-100 (30-50ns). Each column corresponds to a frame, and each row to an interface residue from monomer A or B. Red shades indicate energetically unfavorable contributions, while blue shades indicate stabilizing interactions. The pink and green sidebars designate receptor and ligand monomer residues, respectively (chain A vs chain B).

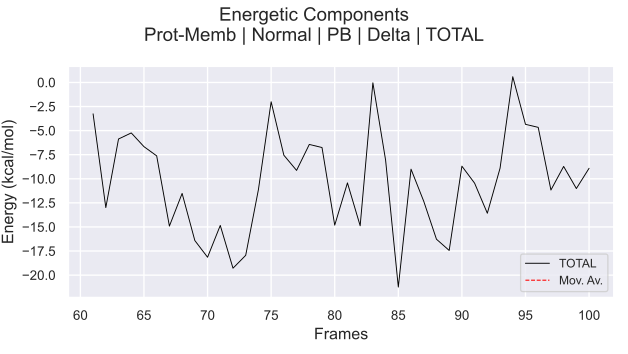

**Figure S12. Total Binding Energy Fluctuations of the VMAT1 - Isoform P Heterodimer Across 50ns full-atom MD simulation in realistic bilayer.** The total MM/PBSA binding energy of the VMAT1 dimer calculated over simulation frames 60-100 (30-50ns). The black line represents the TOTAL binding free energy (kcal/mol), capturing frame-to-frame fluctuations in the overall interaction strength between monomers. The red dashed curve shows a moving average, highlighting underlying energetic trends and smoothing short-timescale variability. Energies include contributions from protein–membrane interactions, van der Waals forces, electrostatics, polar solvation (PB), and nonpolar terms.

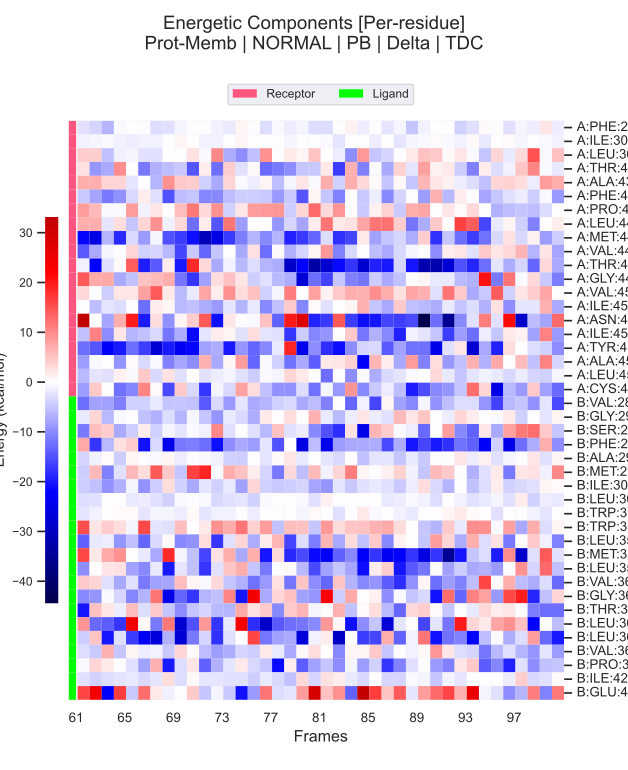

**Figure S13. Per-Residue Binding Energy Decomposition of the VMAT1 - Isoform P Heterodimer across 50ns full-atom MD simulation in realistic bilayer.** Heatmap visualization of residue-wise MM/PBSA energy components (kcal/mol) across simulation frames 60-100 (30-50ns). Each column corresponds to a frame, and each row to an interface residue from monomer A or B. Red shades indicate energetically unfavorable contributions, while blue shades indicate stabilizing interactions. The pink and green sidebars designate receptor and ligand monomer residues, respectively (chain A vs chain B).

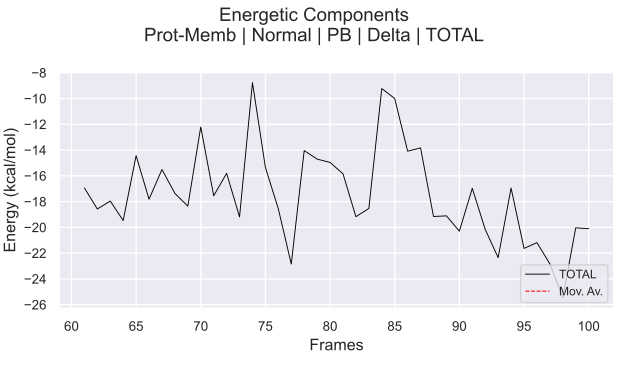

**Figure S14. Total Binding Energy Fluctuations of the Isoform P Homodimer Across 50ns full-atom MD simulation in realistic bilayer.** The total MM/PBSA binding energy of the VMAT1 dimer calculated over simulation frames 60-100 (30-50ns). The black line represents the TOTAL binding free energy (kcal/mol), capturing frame-to-frame fluctuations in the overall interaction strength between monomers. The red dashed curve shows a moving average, highlighting underlying energetic trends and smoothing short-timescale variability. Energies include contributions from protein–membrane interactions, van der Waals forces, electrostatics, polar solvation (PB), and nonpolar terms.

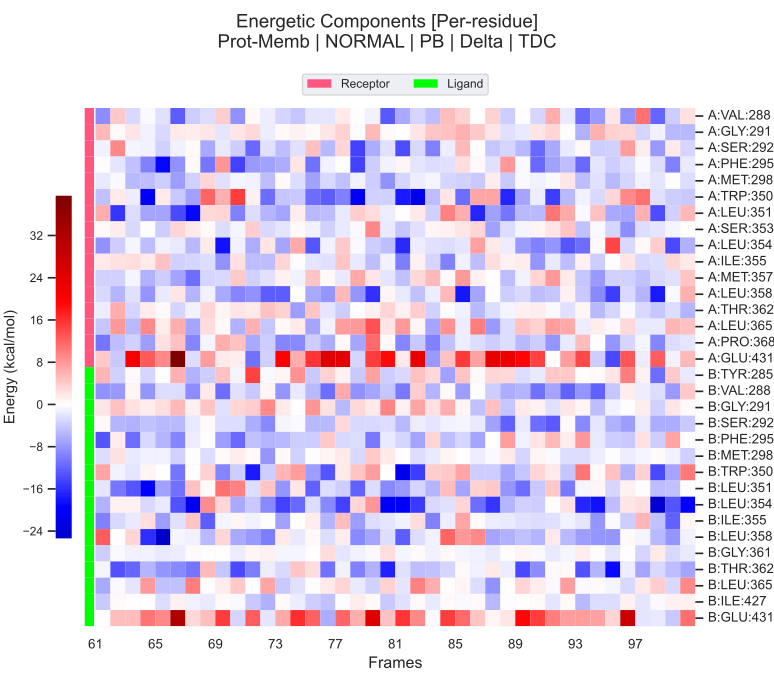

**Figure S15. Per-Residue Binding Energy Decomposition of the Isoform P Homodimer across 50ns full-atom MD simulation in realistic bilayer.** Heatmap visualization of residue-wise MM/PBSA energy components (kcal/mol) across simulation frames 60-100 (30-50ns). Each column corresponds to a frame, and each row to an interface residue from monomer A or B. Red shades indicate energetically unfavorable contributions, while blue shades indicate stabilizing interactions. The pink and green sidebars designate receptor and ligand monomer residues, respectively (chain A vs chain B).

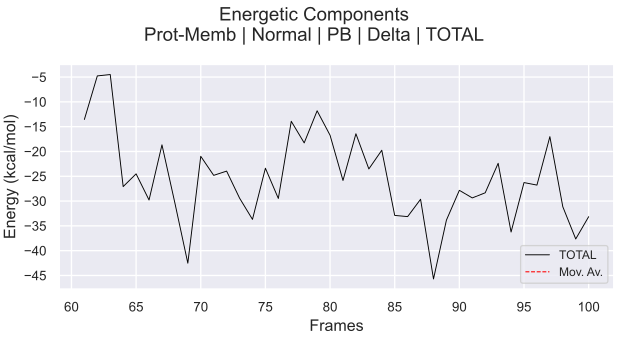

**Figure S16. Total Binding Energy Fluctuations of the Isoform Q Homodimer Across 50ns full-atom MD simulation in realistic bilayer.** The total MM/PBSA binding energy of the VMAT1 dimer calculated over simulation frames 60-100 (30-50ns). The black line represents the TOTAL binding free energy (kcal/mol), capturing frame-to-frame fluctuations in the overall interaction strength between monomers. The red dashed curve shows a moving average, highlighting underlying energetic trends and smoothing short-timescale variability. Energies include contributions from protein–membrane interactions, van der Waals forces, electrostatics, polar solvation (PB), and nonpolar terms.

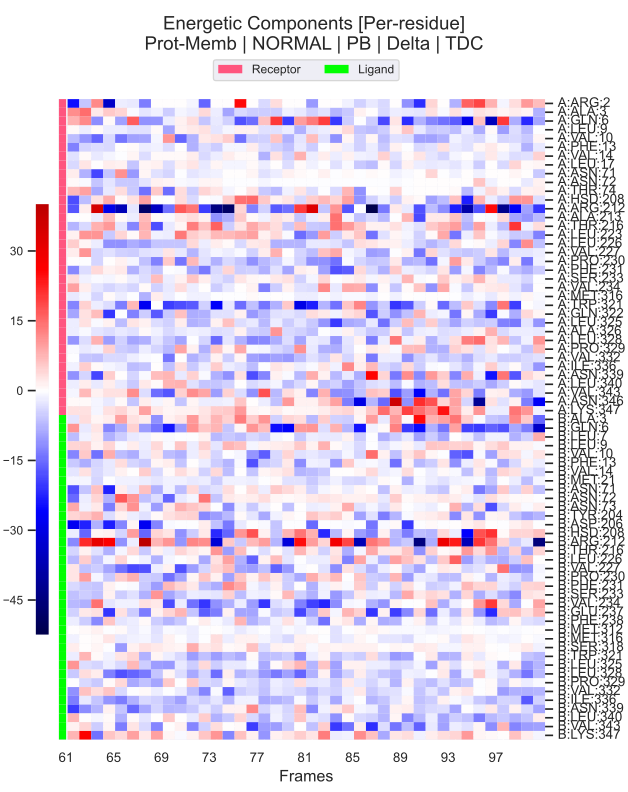

**Figure S17. Per-Residue Binding Energy Decomposition of the Isoform Q Homodimer across 50ns full-atom MD simulation in realistic bilayer.** Heatmap visualization of residue-wise MM/PBSA energy components (kcal/mol) across simulation frames 60-100 (30-50ns). Each column corresponds to a frame, and each row to an interface residue from monomer A or B. Red shades indicate energetically unfavorable contributions, while blue shades indicate stabilizing interactions. The pink and green sidebars designate receptor and ligand monomer residues, respectively (chain A vs chain B).
